# AutoZyme: An Autonomous Agentic Framework to Optimize Bioinformatics Software

**DOI:** 10.64898/2026.06.12.731250

**Authors:** Elliot Xie, Lingxin Cheng, Yujia Cai, Jack Shireman, Christina Kendziorski

## Abstract

Performance bottlenecks in widely used genomics and bioinformatics software present a substantial and growing burden as biological datasets continue to increase in size and number. Relieving these bottlenecks relies largely on expert manual optimization and therefore remains difficult to scale. Here we present AutoZyme, an agentic framework for scientific software optimization. Given a target function, AutoZyme builds benchmarks, identifies bottlenecks, and iteratively tests code changes, retaining only those that improve runtime while preserving output. We evaluated AutoZyme on 45 functions, improving runtime without substantial memory increases in over 95% of cases considered. Across 38 functions from Seurat, Scanpy and related packages in genomics and bioinformatics, AutoZyme reduced runtime by a median of 8.52-fold, with the largest reductions exceeding 676-fold. The optimized functions are distributed through AutoZyme-Library as drop-in replacements for existing analysis pipelines. We also release AutoZyme as a reusable framework for optimizing additional user-specified packages and functions.

## Introduction

Bioinformatics tool development has historically prioritized methodological novelty and statistical accuracy over computational efficiency. This trade-off has become increasingly problematic as single-cell and other high-throughput datasets have grown in scale. We manually audited GitHub and Bioconductor channels for 18 widely used bioinformatics tools and identified 468 genuine complaints citing runtime, excessive memory use, or scaling failures, of which 104 remained unresolved, with a median complaint age of approximately 2 years (Fig. 1a; Methods). The resulting pattern represents a long-tail maintenance problem where many canonical functions remain widely used despite implementation bottlenecks that users repeatedly encounter.

**Fig. 1.**
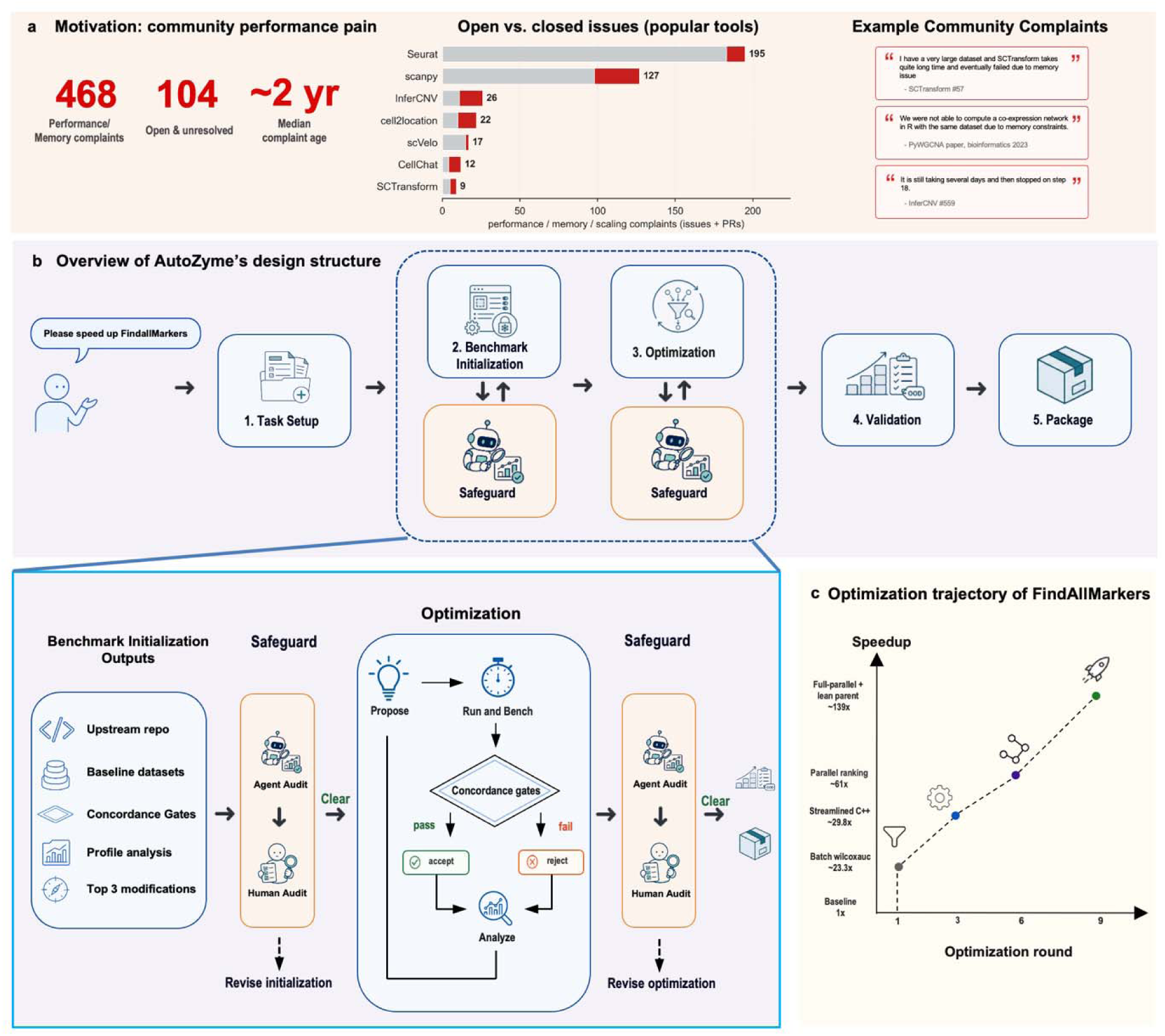
Motivation, design, and FindAllMarkers optimization trajectory. **a,** Community-reported performance, memory, and scaling limitations are common across mature bioinformatics software. A manual audit of GitHub and Bioconductor identified 468 performance or memory complaints, including 104 that remained unresolved and open with a median age of approximately 2 years. The stacked bars show issue status for frequently cited tools: gray denotes closed complaints, and red denotes complaints that remain open and unfixed; labels indicate total complaints per tool. Representative community reports illustrate recurring failure modes including excessive memory use, inability to run analyses at dataset scale, and multi-day runtimes. **b,** Overview of AutoZyme’s modular agentic optimization workflow. The Task Setup agent expects a target repository (e.g. GitHub repo) or function to be optimized and sets up the required file structures. The Benchmark Initialization agent identifies the function to be optimized within the GitHub repo and also identifies benchmarking datasets of varying sizes, runs the function on each dataset to establish baseline results and noise estimates that help determine concordance gates; this agent also profiles the function across datasets to identify bottlenecks and propose three candidate modifications. The Optimization agent considers the three candidates and proposes a candidate for evaluation on the benchmark datasets. If improvement for a candidate patch is greater than expected given noise estimates, the patch is accepted and recorded, and another patch is proposed (iterates for fifty rounds). To reduce the risk of agent hacking, an Auditor agent evaluates results from the candidate after benchmark initialization and after optimization, flagging any suspicions for human audit. Once an optimization passes Auditor review, the optimized patch is moved to the Validation agent for further evaluation. Failed candidates are returned to the Optimization agent (constrained to 30 iterations by default). Once validated, the patches are passed to the Packaging agent for final benchmarking and integration into the AutoZyme-Library. **c,** AutoZyme trajectory for Seurat’s FindAllMarkers. Four accepted patches progressively restructured the computation: batching the Wilcoxon test across clusters (23.3×), composing presto’s C++ primitives directly over a shared transposed matrix (29.8×), running the full per-chunk pipeline in parallel across cores (61×), and folding the parent process’s residual aggregation into the workers themselves (139×; cumulative speedups on the small benchmark).

Existing acceleration strategies, including manual rewrites and targeted compiled kernels, can substantially speed up individual functions [1,2], but require specialized optimization efforts that are difficult to scale. More extensive approaches, including backend replacement libraries such as BPCells[3], accelerate selected workflows through alternative sparse-matrix storage and out-of-core computation, but require substantial tool-specific implementation and may operate through execution paths distinct from the Seurat[4] or Scanpy[5] code paths originally invoked by users. GPU-accelerated frameworks such as the NVIDIA single-cell stack[6] and rapids-singlecell[7] also provide substantial speed improvements for supported operations such as principal component analysis and neighborhood graph construction, but depend on specialized hardware and manually maintained GPU implementations. Moreover, because many bioinformatics workflows operate primarily in CPU-based environments, GPU-based acceleration typically requires additional infrastructure and integration changes extending beyond the optimization itself.

Foundation-model coding agents are another promising approach for automating software optimization[8–11]. However, ad-hoc prompting of coding agents can produce apparent "speedups" when in reality the agent has changed its semantics, evaluated something other than the optimized code, or overfit to the pre-specified benchmarks [12,13] (behavior hereinafter referred to as *agent hacking*). In addition, comprehensively exploring and evaluating optimization options requires extensive human input and interaction.

To address these challenges, we developed AutoZyme, an autonomous agentic framework for scientific software optimization. AutoZyme consists of five main agents: task setup, benchmark initialization, optimization, validation, and packaging. AutoZyme also incorporates an Auditor agent that guards against agent hacking both after benchmark initialization and after optimization. Before optimization begins, AutoZyme establishes the target function, benchmark datasets, upstream reference outputs, output-concordance gates and baseline timing measurements. These components are then fixed before the agent starts modifying code. During optimization, the agent proposes candidate changes with each change executed through the same benchmark pipeline and retained only if it improves runtime while preserving the expected output. Accepted changes are then tested on held-out datasets and matched thread settings to determine whether the speedup persists outside the development benchmark. An independent Auditor agent reviews the benchmark and optimized code to guard against agent hacking. Finally, changes that pass validation are converted into installable patches that preserve the original public API. Across 38 genomics and bioinformatics functions, AutoZyme reduced runtime by a median of 8.52-fold while preserving output concordance and avoiding substantial memory increases in most optimized functions. We release these validated optimizations through AutoZyme-Library, allowing users to accelerate existing CPU-based workflows without rewriting analysis pipelines.

## Results

### AutoZyme is motivated by a long tail of unresolved performance bottlenecks in mature bioinformatics software

We first asked whether performance limitations in mature bioinformatics software represent an isolated implementation problem or a broader maintenance burden. To address this, we manually audited GitHub and Bioconductor for 18 widely used bioinformatics tools and identified 468 issues citing runtime, excessive memory use, or scaling failures; 104 of these remained open and unresolved after a median complaint age of approximately 2 years (Fig. 1a; Methods). These unresolved reports suggest that performance bottlenecks are a persistent feature of bioinformatics software systems requiring scalable approaches to software optimization.

### AutoZyme is a modular, multi-agent large language model (LLM) framework that combines agentic code optimization, benchmarking, and validation to safely optimize scientific software

A schematic illustrating AutoZyme is provided in Fig. 1b, and a summary of each agent is provided below with full details in Methods. The Task Setup agent expects a GitHub repo containing the function to be optimized. It then confirms the user’s intent and establishes a hierarchical directory tree structure to house all files during optimization and packaging. The Benchmark Initialization agent identifies the function to be optimized within the GitHub repo and identifies datasets of varying sizes (small, medium, large) for benchmarking either in the repo, within the AutoZyme framework, supplied by a user, or from public repositories. The agent then runs the function on each dataset with different seeds to establish baseline results and to obtain noise estimates. The noise estimates help determine concordance thresholds for concordance gates, which are established by the agent to preserve output structure, continuous results, and decision-related results. Finally, the Benchmark Initialization agent profiles the function across the benchmark datasets to document how much time each step takes and identifies the top three places in which a code modification may prove worthwhile.

The Optimization agent considers the three candidate modifications suggested by the Benchmark Initialization agent and proposes a specific modification for evaluation on the benchmark datasets (we refer to these modifications as patches). If improvement for a candidate patch is greater than expected given the noise level estimated by the Benchmark Initialization agent, the patch is accepted and recorded, and another patch is proposed. This iterates for 50 rounds. If improvement is consistent across dataset sizes, the optimized function is moved to the Validation agent. The Validation agent identifies two additional independent datasets and runs the optimized function on one thread, four threads, and eight threads to see how performance changes. Failures are returned to the Optimization agent (constrained to 30 iterations by default). Once performance is confirmed to be preserved across dataset sizes and threads, the patches are passed to the Packaging agent for final benchmarking and integration into the AutoZyme-Library.

Finally, AutoZyme incorporates an Auditor agent that guards against agent hacking both after benchmark initialization and after optimization by searching for pre-specified and common, as well as uncommon, hacking patterns.

The AutoZyme agentic workflow is coordinated by *zyme*, a command-line interface (CLI) that codes deterministic steps for repeated, reproducible use. The CLI allows agents to focus on steps that require case-specific reasoning, while *zyme* handles deterministic procedures. Moving these procedures out of prompts shortens the agent context devoted to routine instructions and standardizes steps that would otherwise vary across agent executions; this helps to reduce the likelihood of agent hallucination and/or hacking.

### AutoZyme reduces FindAllMarkers runtime over 100-fold while preserving output concordance

We first applied AutoZyme to Seurat’s FindAllMarkers, a widely used function for identifying cluster-specific marker genes in single-cell RNA-seq data. The Benchmark Initialization agent selected three datasets spanning the typical workload range: a 30K-cell subset of PBMC 68K[14] (small), the full PBMC 68K (medium), and a 120K-cell subset of the PBMC 208K glaucoma[15] dataset (large). Because FindAllMarkers returns a ranked marker table, concordance was measured by preservation of markers: specifically, p-values and log2 fold-changes (Spearman ≥ 0.99), pointwise drift on the numeric columns (99th-percentile absolute difference ≤ 0.02 for p-values ≤ 0.05 and log2FC ≤ 0.01 for detection fractions), and set-level agreement on the significant markers (Jaccard ≥ 0.95, recall ≥ 0.97).

During optimization, AutoZyme accepted a series of patches that progressively restructured FindAllMarkers at four levels of the computation: its per-cluster loop, its boundary with the backend library, its parallel decomposition across cores, and finally the flow of intermediate values between workers. Seurat delegates the Wilcoxon test to a fast C++ backend (presto), but invokes it once per cluster, repeating sparse subsetting, transposition, and gene ranking on every pass. AutoZyme’s first patch reformulated this loop as a single batched evaluation over all clusters, computing per-cluster fold-changes by multiplying the expression matrix against a sparse cluster-indicator matrix. The second patch moved one level below presto’s R interface, calling its C++ primitives directly and threading a single pre-transposed expression matrix through every step. The third patch partitioned the gene set across CPU cores so that each worker carried out the full per-chunk pipeline (ranking, fold-change, filtering, assembly) independently, reducing the parent process to a small aggregation over per-chunk summary. Finally, the fourth patch closed even that gap. Specifically, the agent recognized that each worker, while ranking its own gene chunk, was already computing the same quantities the parent would otherwise aggregate and absorbed the parent’s aggregation into work the workers were already doing. The pipeline became parallel end-to-end (Fig. 1c).

Following optimization, the packaged FindAllMarkers patch was evaluated on the Windows workstation. Runtime fell from ∼1.5 minutes to 1.4 seconds on the small PBMC 68K (71×), from over 11 minutes to 8.1 seconds on the medium PBMC 208K (88×), and from over an hour to 23 seconds on the large Heart 486K (169×) datasets, with all nine concordance metrics holding at their optimal values (Spearman = 1.0, set Jaccard and recall = 1.0, pointwise drift = 0). Peak memory use was also reduced on the same datasets, from 6.6 to 2.8 GB, 28.9 to 11.6 GB, and 70.8 to 29.0 GB, respectively (Fig. 2a).

**Fig. 2.**
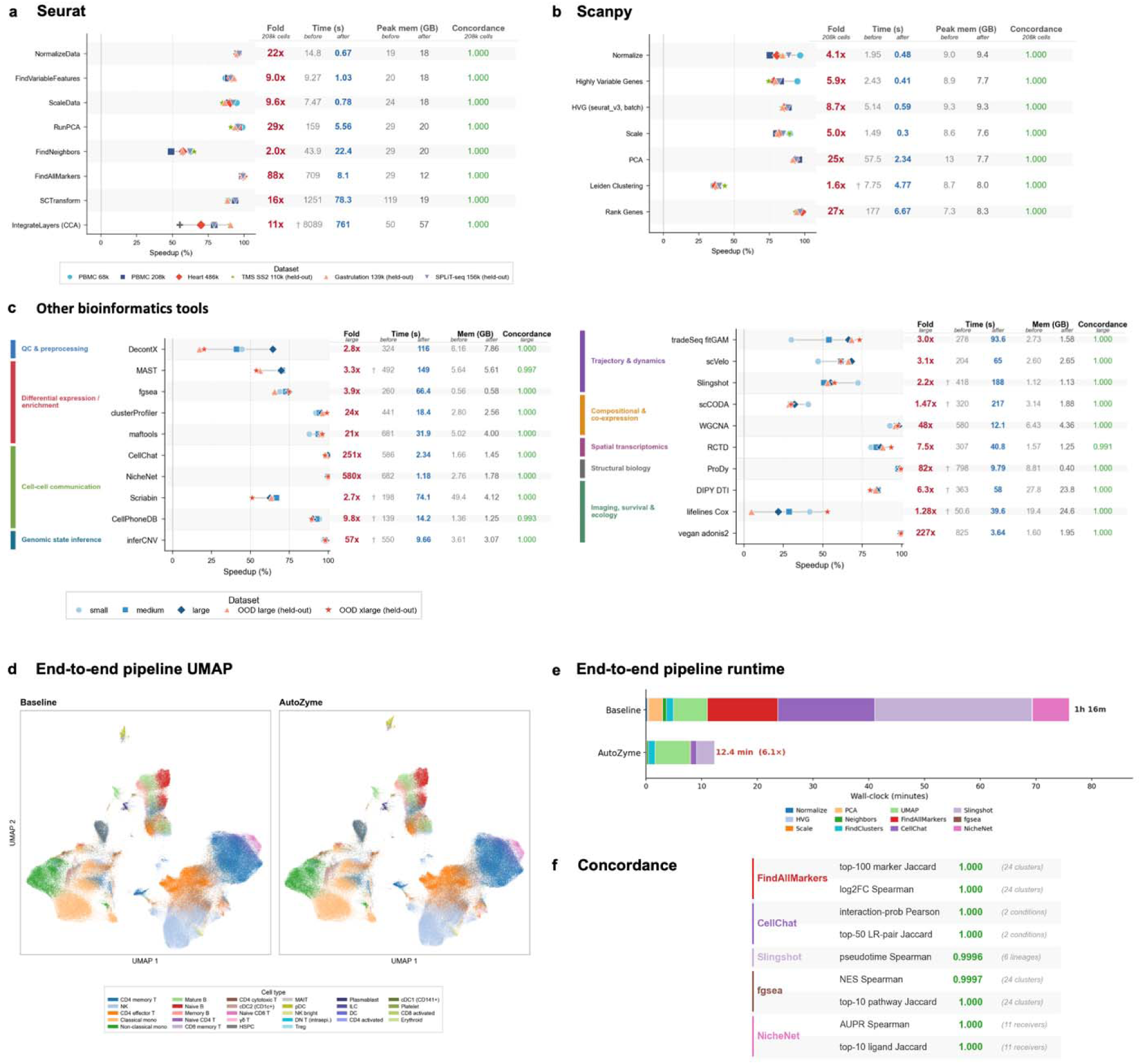
AutoZyme accelerates core single-cell and representative bioinformatics workflows while preserving output concordance. **a,b,** Evaluations of AutoZyme optimized patches for eight Seurat functions **(a)** and seven Scanpy functions **(b)** over 6 benchmark datasets. Points show percentage runtime reduction for individual benchmark datasets; symbols denote PBMC 68K, PBMC 208K, Heart 486K[28], Tabula Muris Senis Smart-seq2 110K[29], gastrulation 139K[30], and SPLiT-seq 156K[31] datasets. Adjacent tables report fold speedup for the PBMC 208K dataset, before/after wall-clock time, before/after peak memory, and task-specific output concordance. Benchmarks were run on a Windows 11 AMD Ryzen 9 7950X workstation with 128 GB DDR5 memory using four threads, values are medians across replicate runs. Across a–c, † marks values reported at one thread; all other values at four threads. Results from a MacBook Pro are shown in Supplementary Data 2. Fold speedup is baseline runtime divided by AutoZyme runtime and Speedup (%) is the percentage of the baseline runtime reduced by AutoZyme, specifically equal to the difference between baseline runtime and AutoZyme runtime divided by baseline runtime; and so a 50% runtime reduction corresponds to a 2-fold change; a 75% runtime reduction to a 4-fold change. The baseline and AutoZyme runtimes were taken under matched data, seeds, parameters, and thread settings. AutoZyme achieved median runtime reductions of 92.4% across Seurat functions and 81.9% across Scanpy functions. **c,** Benchmark summary for representative additional bioinformatics and related scientific-computing tools, grouped by analysis domain, including quality control and preprocessing, differential expression and enrichment, cell–cell communication, genomic state inference, trajectory and dynamics, co-expression and compositional analysis, spatial transcriptomics, structural biology, and imaging, survival, and ecology. **d**, UMAP embeddings from an end-to-end Seurat analysis of the PBMC 208K dataset run with the baseline pipeline (left) and AutoZyme-optimized functions (right). Cells are colored by annotated cell type. Adjacent tables report fold speedup, before/after wall-clock time, before/after peak memory, and output concordance for the large-tier dataset of each tool. **e,** End-to-end wall-clock time for the same Seurat pipeline using baseline or AutoZyme implementations. Stacked bars show the contribution of each analysis step. AutoZyme reduced total runtime from 76 min to 12.4 min, corresponding to a 6.1-fold acceleration. **f,** Downstream concordance between baseline and AutoZyme outputs for FindAllMarkers, CellChat, Slingshot[32], fgsea[33], and NicheNet[34].

### AutoZyme generalizes across genomics and bioinformatics tools while preserving output and memory usage

Having established that AutoZyme could accelerate FindAllMarkers, we next asked whether the same framework could improve performance across a broader set of genomics and bioinformatics tools including those implemented in Seurat as well as other stand-alone packages. As shown in Fig. 2a-c, notable improvements were observed in SCTransform[16], InferCNV[17], WGCNA[18], and CellChat[19] with runtimes reduced 16-, 57-, 48-, and 251-fold and memory usage reduced by 84%, 15%, 32%, and 13%, respectively (per-dataset values in Fig. 2a-c and Supplementary Data 2).

Median runtimes and memory taken over datasets × thread counts × platform configurations show similar results. Full details for each dataset, thread, and configuration are shown in Supplementary Data 2 and summarized in Supplementary Note 10 and Supplementary Fig. 5. As detailed there, across 38 genomics and bioinformatics functions, AutoZyme reduced runtime by a median of 8.52-fold while preserving output concordance. Memory usage was within 15% of the unoptimized baseline in 97.4% of optimized functions (37 of 38 functions).

To assess the extent to which per-function gains translated into workflow improvements, we chained AutoZyme-optimized functions to conduct a full Seurat analysis of the PBMC 208K data. The analysis on an AMD Ryzen 9 7950X workstation using 128 GB DDR5 and 4 threads took 76 minutes whereas the AutoZyme-optimized workflow completed in 12.4 min, corresponding to a 6.1-fold reduction (Fig. 2e). Analyses preserved concordance across all five tasks, with Jaccard and Spearman/Pearson coefficients of at least 0.999 (Fig. 2a-d, f).

As AutoZyme is not specific to genomics and bioinformatics, we also applied the framework to 7 packages widely used in other scientific domains including astronomy (astropy), seismology (ObsPy), satellite remote sensing (sarsen), climate science (xclim), computational fluid dynamics (FiPy), molecular dynamics (MDAnalysis), and statistical inference (statsmodels). Across these packages, AutoZyme achieved a median per-task speedup of 9-fold (range 3.5–424-fold), with concurrent memory reductions in 6 of 7 cases. Per-task performance summaries are provided in Supplementary Note 11, Supplementary Fig. 2 and Supplementary Data 2 (Non-bio tab).

### AutoZyme-Library provides drop-in acceleration for 45 functions

Successful optimizations of 45 functions were packaged for reuse in AutoZyme-Library as validated patches registered against the original public APIs. The current library spans core Seurat and Scanpy routines, commonly used bioinformatics tools including CellChat, InferCNV, WGCNA, and seven selected packages from other scientific fields (Fig. 3a).

**Fig. 3.**
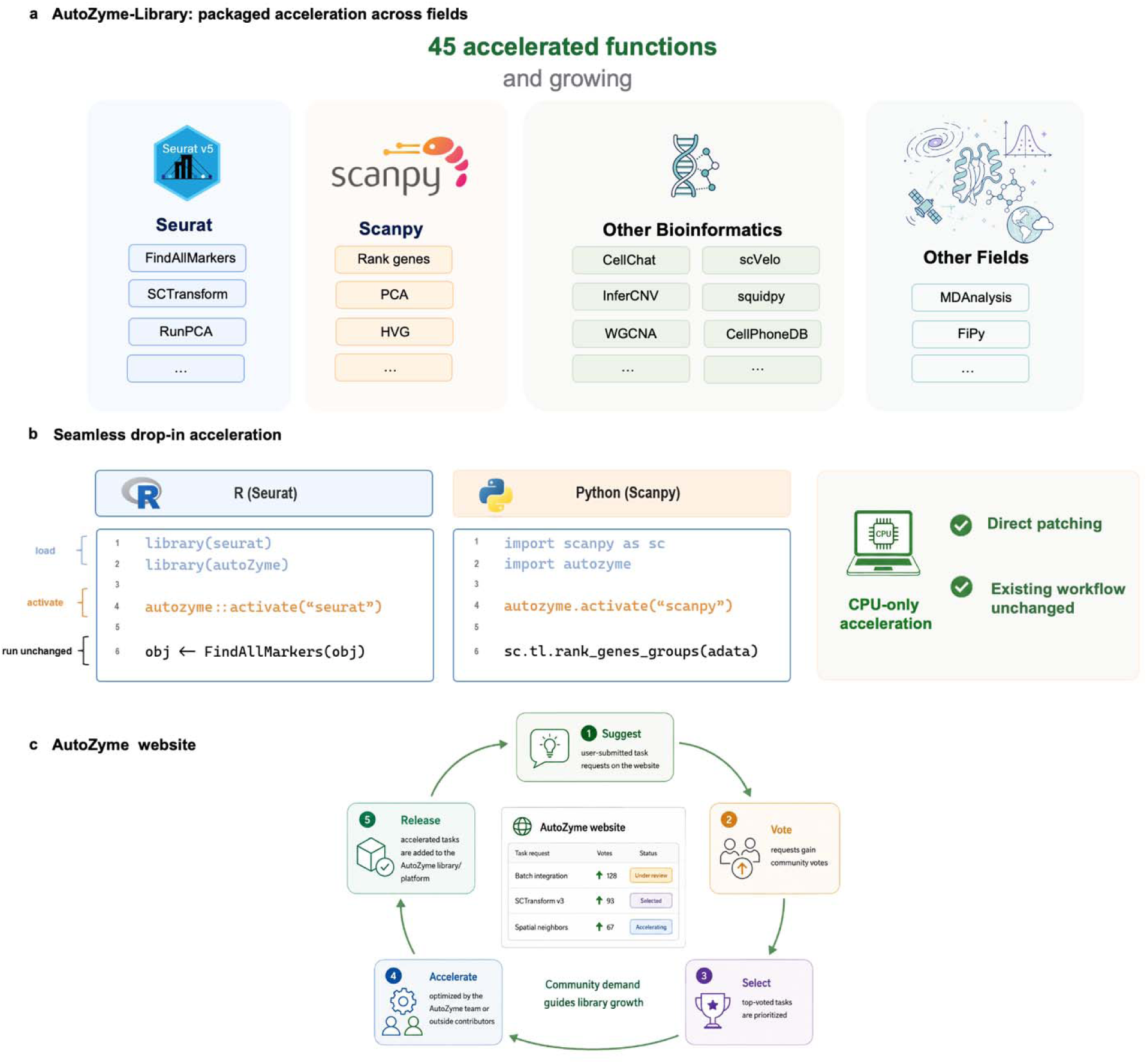
AutoZyme-Library packages validated optimizations as drop-in accelerators for existing scientific workflows. **a,** Scope of AutoZyme-Library. Validated AutoZyme optimizations are distributed as a growing collection of 45 optimized functions spanning core Seurat routines, Scanpy workflows, additional bioinformatics packages and selected non-biological scientific-computing tools. Boxes show representative supported functions or packages. **b,** Drop-in activation model. In R or Python, users load AutoZyme-Library, activate the relevant package patch and then call the original upstream function names, such as FindAllMarkers or sc.tl.rank_genes_groups, without modifying the surrounding analysis workflow. Activated patches directly dispatch the targeted calls to CPU-only optimized implementations while preserving the public API. **c,** Community-guided expansion of the library. The AutoZyme website (autozyme.com) provides a request and prioritization loop in which users suggest functions or packages for acceleration, community votes identify high-impact bottlenecks, selected tasks are optimized and validated, and successful accelerations are released back into AutoZyme-Library.

As AutoZyme-Library preserves the original public APIs and surrounding workflow structure, optimized implementations can be introduced without modifying existing analysis pipelines. Specifically, in R and Python, users load AutoZyme-Library, activate the relevant package patch, and then call the same upstream function names as before, such as FindAllMarkers or sc.tl.rank_genes_groups (Fig. 3b).

To support continued expansion, we also built the AutoZyme website as a community request and prioritization portal. Users can nominate functions or packages that require acceleration, vote on high-impact bottlenecks that they would like to see addressed, and track selected requests as they move through optimization, validation and release into AutoZyme-Library (Fig. 3c). In this way, AutoZyme-Library provides optimizations that can be easily integrated into existing scientific workflows and extended according to community demand.

### AutoZyme discovers recurrent structural mechanisms rather than isolated one-off patches

We next asked whether AutoZyme’s accepted optimizations reflected recurring performance mechanisms or isolated package-specific edits. Across the AutoZyme-Library, accepted patches grouped into five broad mechanism families: redundant work removal and reuse, execution path and representation changes, kernel engineering, exact algorithmic reformulation, and controlled numerical relaxation (Fig. 4a).

**Fig. 4.**
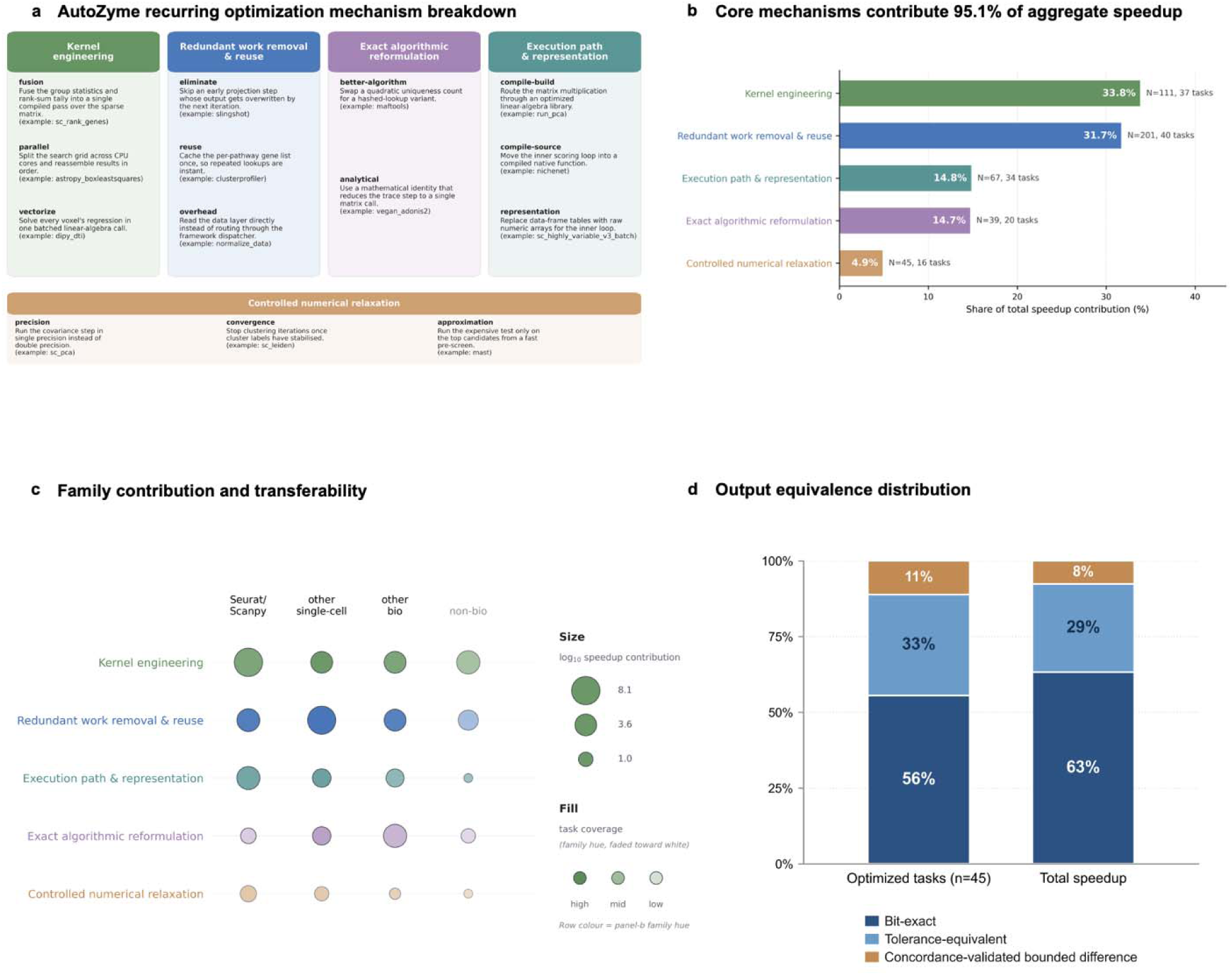
AutoZyme discovers recurrent mechanisms of acceleration. **a,** AutoZyme optimizations were classified into kernel engineering, redundant work removal and reuse, execution path and representation changes, exact algorithmic reformulation, and controlled numerical relaxation, with representative examples shown for each class. **b,** Bars show the summed contribution across patches within each mechanism, and in-bar percentages denote each mechanism’s share of total attributed speedup. The number of patches assigned to each family is denoted by *N*, followed by the number of distinct optimized functions containing at least one such patch. **c,** Mechanism-family contribution and transferability across task groups. Circle area shows the summed log10 speedup contribution from accepted patches in each mechanism–task-group pair; fill intensity shows task coverage, defined as the fraction of tasks in that group with at least one accepted patch from that family. Row colors match the mechanism-family palette in b. **d,** Output-equivalence distribution of final optimized function endpoints. The left bar is normalized by the number of optimized functions (*n* = 45); the right bar is normalized by aggregate speedup contribution. Colors indicate bit-exact, tolerance-equivalent and concordance-validated bounded-difference outputs; percentages are normalized separately within each bar.

Redundant computation removal and reuse accounted for the most patches (201 edits), and the underlying cases were often simple once identified. Beyond this baseline cleanup, kernel engineering was the largest single contributor to aggregate speedup at 33.8%, reflecting genuine systems work such as fused rank-sum passes over sparse matrices in sc_rank_genes; execution path and representation changes added another 14.8% by routing inner loops through compiled code or numerical arrays. Exact algorithmic reformulation contributed a further 14.7% of speedup, in cases where the agent rewrote the underlying computation while keeping the output bit-exact, including a linear-algebra identity that collapses per-term trace. Controlled numerical relaxation, by contrast, contributed only 4.9%, indicating that AutoZyme accelerates existing software primarily by finding more efficient implementations of the same computation rather than by changing output (Supplementary Note 6, Supplementary Table 1 and Supplementary Fig. 3 provide examples for each type of patch, and three representative patches are worked through in detail in Supplementary Note 7). Consistent with this, 40 of 45 tasks (89%) match the original output exactly or within 0.6% drift and deliver 92% of the total speedup; the remaining 5 tasks were also within their concordance gates, with drift of 1.5% to 5% from the baseline (Fig. 4b, d). The same mechanism families recurred across Seurat, Scanpy, broader bioinformatics packages, and packages in other areas (Fig. 4c).

## Discussion

Performance optimization is an essential part of maintaining scientific software that typically requires specialized engineering effort, detailed knowledge of the target package, or hardware and infrastructure that may not be widely available. LLMs are an attractive approach to software optimization, but ad-hoc applications are prone to benchmark overfitting and agent hacking. To address these limitations, we developed AutoZyme, a multi-agent LLM framework designed for automated scientific software optimization. AutoZyme’s multi-agent structure consists of Task Setup, Benchmark Initialization, Optimization, Validation and Packaging agents. Comprehensive quality control is provided through function-specific concordance gates, benchmarking, held-out validation datasets, multi-threaded scaling tests and an additional Auditor agent that screens specifically for agent hacking. In addition to producing optimized code, AutoZyme records the full optimization history, including accepted patches as well as benchmark and concordance test results. This allows users and maintainers to inspect how each optimization was obtained and the extent to which each preserves output.

AutoZyme exemplifies how CLI-assisted multi-agent LLM frameworks can move beyond ad-hoc prompting to perform structured software engineering that leads to effective optimizations. Across 38 genomics and bioinformatics functions, AutoZyme reduced runtime by a median of 8.52-fold, with the largest gains exceeding 676-fold, while preserving task-specific output concordance and avoiding substantial memory increases in most optimized functions. The results shown here used AutoZyme driven by Claude Code with Opus 4.7, but the framework can also be run with other coding agents. Independent Claude Code, Codex and Cursor runs can also identify complementary optimizations, and in two tasks, compatible changes were fused to produce additional gains (Supplementary Note 9 and Supplementary Table 3).

A central design choice underlying AutoZyme is *zyme*, a CLI system that conducts the fixed and repeatable parts of the optimization. This separates the agents’ primary role which is to reason about how to improve the implementation from deterministic tasks such as running benchmarks, recording runtime and memory, comparing outputs with the frozen reference, managing accept-or-reject decisions and restoring rejected code. The same design also makes optimization results auditable, because each accepted patch remains linked to the code changes, benchmark evidence and concordance measurements that supported it. Finally, because critical steps are executed through *zyme* rather than left to the agent’s discretion, AutoZyme can apply checks during optimization and reduce the risk of invalid speedups caused by changed benchmarks, time-window hoisting, or other forms of agent hacking.

To make validated optimizations usable in routine analysis, we distribute successful patches through AutoZyme-Library. The library provides a curated set of optimizations for 45 functions that preserve the original public APIs, allowing users to run optimized implementations within existing CPU-based workflows without rewriting pipelines or requiring specialized hardware. Users can also run the AutoZyme framework independently to optimize additional functions and maintain local AutoZyme-Library instances. Finally, to support community-driven expansion, the AutoZyme website allows users to nominate packages or functions for optimization and/or vote on high-impact bottlenecks for incorporation into AutoZyme-Library.

In summary, AutoZyme provides an optimization tool for bioinformatics software development. By identifying performance-improving patches, preserving output, guarding against agent hacking, and packaging successful optimizations for downstream use, AutoZyme provides optimizations that result in substantially reduced runtime (without substantial increases in memory) across widely used genomics and bioinformatics tools. The AutoZyme framework as well as the optimizations contained within the AutoZyme-Library are expected to prove useful to a wide range of investigators.

## Methods

### Overview

AutoZyme is a multi-agent LLM framework consisting of five interconnected agents: Task Setup, Benchmark Initialization, Optimization, Validation and Packaging. An independent Auditor agent is also included to guard against agent hacking. Each agent is described in detail below, with full prompts for each agent provided in Supplementary Note 12 and Supplementary Fig. 4. The AutoZyme workflow is coordinated by *zyme*, a command-line interface (CLI) that codes deterministic steps for repeated, reproducible use. The CLI allows agents to focus on steps that require case-specific reasoning, such as deciding the next modification to test or assessing whether an optimized task reflects agent hacking. In contrast, *zyme* handles fixed procedures including profiling the target function, executing benchmark runs, recording timing, memory and concordance metrics, and maintaining task state. The full subcommand reference and task-directory layout are provided in Supplementary Note 3. Moving these procedures out of prompts shortens the agent context devoted to routine instructions and standardizes steps that would otherwise vary across agent executions; this helps to reduce the likelihood of agent hallucination and/or hacking. In addition, a CLI interface reduces token use. For the Optimization agent specifically, because each critical action of the agent (e.g. run the benchmark, accept/reject candidate modification) is routed through the CLI, *zyme* can also apply checks before a candidate is accepted at every iteration round to prevent agent hacking and can return structured feedback when the agent deviates from the expected optimization trajectory.

### Task setup

The Task Setup agent takes a user-specified GitHub repo and target function as input. After confirming the target, it runs *zyme init* to create a version-controlled task directory containing the templates and metadata required for executing and evaluating the function of interest. User-supplied dataset paths are checked and stored; when no path is provided, benchmark datasets are selected during Benchmark Initialization.

### Benchmark Initialization

The Benchmark Initialization agent identifies the function to be optimized within the GitHub repo, checks that the function version at the GitHub repo matches the installed version if available, and records the exact argument settings used for benchmarking. The agent then identifies datasets of varying sizes (small, medium, large) for benchmarking either in the repo, supplied by a user, or downloaded from public sources. Using the templates created during Task Setup, the agent writes three scripts: a reference script that runs the upstream implementation, a candidate script that runs the current optimized implementation, and an evaluation script that compares candidate outputs with reference outputs. These scripts read dataset-specific input and output paths, allowing the same benchmark pipeline to run across all selected datasets. The agent then runs *zyme baseline* to evaluate the function on each dataset with different seeds to establish baseline results and to obtain noise estimates. The noise estimates help determine concordance thresholds for concordance gates, which are established by the agent to preserve output structure, continuous results, and decision-related results.

### Concordance Gates

For each task, AutoZyme defines three classes of concordance gates before optimization to preserve output structure, continuous results, and decision-related results. The output structure gate preserves the documented user-facing return slots of the function, including the presence, shape and non-degeneracy of each output. This check is executed automatically by *zyme* during baseline and optimization runs. The gates associated with preserving continuous results and decision-related results are implemented as task-specific metrics in the evaluation script, with thresholds fixed before optimization.

Continuous-output gates are used for scores, probabilities, fitted values, weights, embeddings, predicted expression values and other numerical outputs. These gates pair a correlation metric with a pointwise drift metric. Spearman correlation is used when downstream interpretation depends mainly on rank, whereas Pearson correlation is used when the magnitude of the values is the relevant quantity. Pointwise drift is usually measured by the 99^th^ percentile absolute difference rather than the maximum absolute difference, reducing sensitivity to a single extreme value. Compositional outputs were evaluated using per-row L1 drift together with a rank or magnitude correlation.

Decision-related gates are used when outputs define user-visible calls, such as cluster assignments, selected gene sets, top-ranked markers or pathway calls. These gates use agreement metrics matched to the decision type, including label agreement, selected-set Jaccard similarity, recall, top-k overlap and absolute call-rate difference. When a continuous score determines a downstream decision, AutoZyme evaluates both the score and the resulting decision.

For deterministic tasks, gate thresholds are fixed before optimization. For stochastic tasks, AutoZyme estimates variability in the function output across seeds and defines effective thresholds from an absolute floor and a multiplier of the estimated noise. This allows ordinary stochastic variation in the original implementation while rejecting optimization-induced drift outside the expected noise level. Task-specific metrics and thresholds are listed in Supplementary Data 1.

### Profiling

Once concordance gates are established by the Benchmark Initialization agent, the agent profiles the function on the small benchmark dataset to document how much time each step takes and identifies the top three places in which a code modification may prove worthwhile. Profiling is carried out through *zyme profile*. Backend-specific details are described in Supplementary Note 3 under profile subcommand.

Once concordance gates are established and profiling is complete, the proposed concordance gates, metadata, and benchmark results are audited by the Auditor agent (see Auditor agent below). The gates, scripts, and benchmark datasets are then frozen before optimization. The Optimization agent does not have permission to modify these components.

### Optimization

The Optimization agent improves the target function iteratively. At the start of optimization, the agent reads the task metadata, evaluation script, reference script, candidate script, recent benchmark results and the top three optimization candidates identified by the Benchmark Initialization agent. Based on this information, the Optimization agent proposes a candidate modification to the current implementation.

The candidate is then executed through the task pipeline using *zyme run*. Each run automatically records the agent’s hypothesis, dataset tier, thread setting, runtime, peak memory and concordance-gate metric values. After reviewing these results, the agent must explicitly accept or reject the candidate modification. Accepted modifications advance to another iteration; rejections are logged together with the evidence against them, and the task folder is restored to the current-best implementation, so that rejected code does not contaminate subsequent measurements. This accept-or-reject cycle is repeated as the agent searches for additional improvements; per-task rejection counts at each stage are reported in Supplementary Data 3 and Supplementary Fig. 6 shows accepted results are concordance-preserving, not merely the fastest.

A candidate modification is accepted when the implementation passes the frozen concordance gates, improves runtime relative to the relevant current-best implementation by more than the expected measurement noise, and avoids a significant increase in memory. Borderline timing changes are remeasured with replicated reruns before acceptance.

Optimization usually proceeds on the small benchmark dataset to allow rapid iteration, with evaluation on the medium and large benchmark datasets primarily conducted via the Validation agent. However, performance on the medium and large datasets is evaluated via the Optimization agent when a proposed modification could alter scale-dependent behavior, when the bottleneck is absent at small scale, when cluster-sensitive outputs are involved since the number of clusters may change with dataset size, or when a preliminary small-tier improvement requires scale confirmation.

We also enforce an Amdahl-first parallelism discipline for the optimization agent[20]. Specifically, the agent recognizes that serial-path improvements are preferred before adding new parallelism. If a candidate changes an upstream-exposed parallelism setting, the corresponding upstream baseline will be measured at the same thread setting.

### Validation

After optimization converges and the accepted patch passes Auditor review (see below), it is transferred to the Validation agent. This step tests whether the optimization still holds outside the development setting: on datasets from different biological contexts or modalities, on inputs larger than those used during optimization, and at thread counts that were not used during development. In addition to the benchmark datasets of varying sizes (small, medium, and large) identified by the Benchmark Initialization agent, the Validation agent defines two additional independent datasets. A large out-of-distribution dataset (OOD-large) is similar in scale to the large benchmark dataset but changes the biological context, modality or execution path. OOD-xlarge is larger than any benchmark dataset considered and is used to test whether the patch still works when input size and wall-clock time increase substantially. These held-out datasets are evaluated across a thread-count matrix, typically 1, 4 and 8 threads. Before running this matrix, AutoZyme determines how the baseline implementation should be measured at each thread count. If the optimized implementation uses threading, the baseline must also be defined for the same thread counts so that speedups are not inflated by an unmatched comparison. For each dataset and thread count, AutoZyme runs the optimized and baseline implementations with matched inputs and parameters. If the baseline fails, exceeds memory or cannot be measured, that setting is marked as unmeasurable and is not used to calculate speedup; the full registry of dataset exclusions, fallbacks and not-applicable tiers is given in Supplementary Note 5.

The measured results are then judged in two ways. First, AutoZyme asks whether the speedup observed during development (calculated on the small, medium, and large benchmark datasets) is preserved on the OOD datasets. For each OOD dataset at a given thread count, the Validation agent compares the development speedup to the speedup calculated on the OOD datasets using the same thread count. If the speedup in the OOD datasets is more than fivefold lower than the development speedup, the patch fails validation. Smaller losses are treated as warnings: 1.5- to 5-fold for OOD-large and 2- to 5-fold for OOD-xlarge. A warning triggers one diagnostic round and is allowed only when the loss can be explained by a documented hardware limit or input-scale limit rather than by a fixable problem in the optimized code. Second, AutoZyme checks whether the threaded speedups are plausible, for tasks where both the baseline and the optimized code expose a parallel path. At *N* threads, the speedup must be at least as large as the speedup at one thread, because adding threads should not make the optimized implementation relatively worse. It also must not exceed 1.5 × *N* times the one-thread speedup. Larger gains are treated as suspicious because they often indicate that the one-thread optimized path is broken or unfairly slow. Either violation fails validation.

Any validation failure, or any warning without a documented cause, opens a validation repair loop capped at 30 rounds. If the failure cannot be fixed within that budget, the patch is kept as long as it still delivers a meaningful speedup and is otherwise rolled back or excluded from packaged results. A patch passes validation only when all measurable dataset and thread-count settings pass the concordance checks, preserve enough of the development speedup on held-out data, show plausible threaded behavior, and pass a final stability check in which the medium and large benchmarks are re-run three times at each thread count to confirm the optimized results are reproducible.

### Packaging

After a candidate optimization passes validation, the Packaging agent converts the implementation from the task folder into an installable patch in the AutoZyme R or Python package. Before final benchmarking, the Packaging agent records which public API call is supported, which inputs and parameter settings use the optimized implementation, and which cases must fall back to the original upstream implementation. The optimized path cannot cover behavior that was not included in the task signature or explicitly tested during validation. This prevents task-specific accelerations from silently intercepting untested branches of the upstream API. After installation of the patch, AutoZyme runs *zyme package preflight*. This command checks the installed patch through the public API, including a minimal test run; output compatibility and portability issues are also checked. If preflight reports platform-specific behavior, the Packaging agent either replaces it with a portable implementation or adds an explicit fallback; *zyme package preflight* is rerun until the patch passes. When only one operating system is available locally, passing preflight is used as the package-level correctness gate, while speed on other operating systems is recorded as pending unless measured separately. Final measurement is performed with *zyme attest*. Outputs are evaluated using the same frozen correctness metrics used during validation. Unless stated otherwise, reported speedups are therefore package-level speedups from installed patches invoked through the user-facing API.

### Auditor agent and additional measures to prevent agent hacking

We use agent hacking to describe cases in which an agent obtains an apparent speedup by changing the benchmark, bypassing the intended code path, narrowing the tested behavior or otherwise producing a result that cannot be shipped as a valid acceleration. AutoZyme uses two complementary safeguards against these failures.

The first is built into the *zyme* CLI, which checks four aspects of each optimization round. First, it verifies that the optimized code actually ran. A candidate run is invalid if the expected activation marker is missing, if the upstream binding was not changed, or if the run used a stale binding or unsupported upstream version. Second, it protects the frozen correctness gate. AutoZyme parses only metrics declared in the frozen task metadata, and it flags edits to the evaluation script, framework or other out-of-scope files instead of treating them as valid optimization progress. Third, the *zyme* CLI keeps the optimization trajectory recoverable. Each round is recorded through version-controlled accept, reject and rollback steps, so rejected code is removed before the next round and accepted rounds can be demoted if later evidence invalidates them. Fourth, it protects measurement integrity. AutoZyme rejects zero-change rounds, measures baselines separately for each dataset tier and thread count, fixes thread-related environment variables, repeats measurements when timing changes fall within the noise estimates, and records crashes or out-of-memory events as failures rather than omissions.

The second safeguard is an independent Auditor agent, invoked through *zyme validate*, which reviews cases that require task-specific judgment. Specifically, the Auditor agent runs after benchmark initialization and again after optimization. It is invoked through *zyme validate* with read-only access to the initialized task folder. It is required to inspect the candidate implementation, pipeline, evaluator, task metadata, recent profiling and benchmark evidence, and task memory; it may also review other task-local files when needed. The Auditor cannot modify code, metadata or framework files; its only output is a structured findings report of agent-hacking mechanisms. High-severity findings require human review before the candidate can move to validation or packaging. Lower-severity findings are recorded for trend monitoring but do not block the round. The severity definitions and Auditor mechanism categories are provided in Supplementary Note 8, Supplementary Figs. 7 and 8 and Supplementary Table 4; per-task Auditor findings are retained in the version-controlled task provenance logs (.zyme/audit.jsonl) in the AutoZyme repository.

### AutoZyme applications

For a given function, AutoZyme was run on a MacBook Pro with an Apple M3 Max processor and 36 GB RAM to optimize the function. Once the optimization was obtained, it was evaluated for both speed and memory on the MacBook Pro and on a Windows 11 workstation with an AMD Ryzen 9 7950X processor, 16 cores, 128 GB DDR5 memory, and 32 threads. Evaluations were obtained at varying thread counts as specified in the text. Results associated with speed and memory reported in the text are from Windows runs, with results from MacBook Pro runs shown in Supplementary Data 2. The exception to this is Figure 1c since it shows speed differences obtained during optimization before a particular patch is determined.

### Benchmarking protocol

Benchmarks were run on two systems: a MacBook Pro with an Apple M3 Max processor and 36 GB RAM, and a Windows 11 workstation with an AMD Ryzen 9 7950X processor, 16 cores, 32 threads and 128 GB DDR5 memory. Evaluations were obtained at varying thread counts as specified in the text. Package installation, patch activation, optimization-loop benchmarking and package-level concordance verification were tested on both systems when supported by the task dependencies. Baseline and optimized measurements used the same input data, random seeds, thread count and user-level parameters, with only the upstream implementation versus the AutoZyme-patched implementation varied. Unless otherwise stated, the benchmark values reported in the manuscript were measured on the Windows workstation using four threads and are reported as the median across replicate package-level runs; the per-task replicate count is reported in Supplementary Data 2. Windows was chosen as Apple M3 runs failed on some of the largest datasets; Apple M3 thread settings are described in Supplementary Note 4, and available platform-specific cross-domain measurements are shown in Supplementary Fig. 2; Seurat and Scanpy Apple M3 results are in Supplementary Data 2 (Seurat and Scanpy tabs) and Supplementary Fig. 5.

Runtime was measured as wall-clock time for the timed public API call. Peak memory was measured as peak resident set size during the measured run. Imports, data loading, object construction and output serialization were excluded from the timed region unless they were part of the target public API call. For every task, Supplementary Data 1 reports the target function, repository and package version, the programming language, the development and held-out dataset tiers, the concordance gate metrics with their thresholds, and the final concordance class. Concordance classes were reported as bit-exact, tolerance or bounded. Seurat and Scanpy benchmark suites are additionally summarized in Supplementary Data 2 (Seurat and Scanpy tabs). Package versions, operating-system details, environment variables, BLAS/OpenMP settings and task-specific environment notes are reported in Supplementary Note 2 and Supplementary Data 1; dataset sources, accessions and the Hugging Face mirror are provided in Supplementary Note 1 and Supplementary Data 4.

Evaluation of AutoZyme optimized patches for eight Seurat functions and seven Scanpy functions across six benchmark datasets are shown in Figure 2. Since IntegrateLayers (CCA) requires multi-batch input, it is benchmarked on six multi-batch datasets: four of the six core datasets together with the IFN-β-stimulated PBMC dataset and the multimodal CITE-seq reference. Sizes and batch counts are shown in Supplementary Data 4; per-dataset results in Supplementary Data 2.

### Mechanism annotation and speedup attribution

Optimized patches are classified as redundant work removal and reuse, execution path and representation changes, kernel engineering, exact algorithmic reformulation, and controlled numerical relaxation. A rubric was developed by reviewing accepted patches across the benchmark suite. After the rubric was fixed, LLM agents labeled each accepted round using the round summary, commit description and task memory. The authors then audited all labels and corrected them as needed. The full labeling rubric is provided in Supplementary Table 1, and the detailed speedup attribution in Supplementary Fig. 3.

Redundant work removal and reuse covers changes that removed repeated computation, cached intermediate results, reused data structures or avoided computing quantities that were later overwritten. Execution path and representation changes cover edits that use a more efficient existing backend, change sparse or dense representations, reduce conversion overhead or select a better dispatch path while preserving the same user-facing output. Kernel engineering covers fused loops, compiled kernels, vectorized inner loops, sparse-matrix kernels, memory-locality improvements and other low-level rewrites. Exact algorithmic reformulation covers mathematically equivalent changes that altered the computational form of the method while preserving bit-exact or near-bit output. Controlled numerical relaxation covers bounded approximations or tolerance changes that remain within the frozen concordance gate but are not claimed to be strictly identical to the upstream output. Speedup contributions were calculated along the accepted optimization trajectory. For each accepted candidate modification, the change in log speedup relative to the previous implementation is attributed to that modification’s classification.

### Community audit of performance complaints

To assess whether performance limitations in mature bioinformatics software represent a broad maintenance burden, we audited community-reported complaints about speed, memory use and scaling across 18 widely used single-cell and bioinformatics tools. For tools with comparable GitHub issue tracking, we searched issues and pull requests on 29 April 2026 using 12 plain-text terms related to speed, memory and scale: slow, memory, OOM, out of memory, takes too long, performance, runtime, crash, large dataset, too slow, RAM and hours. Search results were deduplicated by issue or pull-request number. For WGCNA, which did not have comparable issue tracking, we manually curated performance-related reports from Bioconductor Support, Biostars, maintainer documentation and downstream-tool publications.

The GitHub search returned 1,056 candidate items from 18,913 total issues and pull requests. Candidates were first triaged by a rule-based classifier as genuine performance reports, false positives or borderline cases. LLM-assisted batch review was then used to remove remaining non-performance reports, and each retained thread was reviewed to determine whether a fix had shipped by the audit date. After this review, we retained 456 genuine GitHub complaints and 12 manually curated WGCNA architectural entries, for 468 performance, memory or scaling complaints in total. Of these, 104 remained open or unresolved at the time of audit, comprising 92 GitHub complaints and all 12 WGCNA architectural entries. The median age of unresolved complaints was approximately two years. Per-tool complaint counts and the age distribution are shown in Supplementary Fig. 1, and the full per-item audit ledger in Supplementary Data 5.

### Cross-domain applications

In addition to software widely used in genomics and bioinformatics, we applied the AutoZyme framework to functions used in astronomy, seismology, remote sensing, climate science, computational fluid dynamics, molecular dynamics and applied statistics. These included *astropy.BoxLeastSquares*[21], a periodogram search for exoplanet transit detection; *obspy.Stream*[22] for bandpass filtering; *sarsen*[23] for synthetic-aperture-radar terrain correction; *xclim*[24] for season-length climate-index computation; *fipy*[25], a finite-volume PDE solver; *MDAnalysis*[26] *RMSD* for molecular-dynamics trajectory analysis and *statsmodels*[27] GLM for Poisson/log-link fitting. These cross-domain tasks use the same AutoZyme framework, with minor changes to domain-specific prompts.

## Data availability

All datasets used in this study are publicly available. Benchmark inputs were derived from the following public sources. The 68k PBMC dataset was obtained from 10x Genomics (https://www.10xgenomics.com/datasets) [14], and the ∼208,000-cell PBMC glaucoma atlas from CZ CELLxGENE (https://cellxgene.cziscience.com/) under Gene Expression Omnibus (GEO) accession GSE268936 [15]. The IFN-β-stimulated PBMC dataset was obtained through SeuratData (ifnb; GEO GSE96583) [35], and the multimodal PBMC CITE-seq reference from the Seurat v4 reference of Hao et al. [36]. The adult human heart atlas was obtained from CZ CELLxGENE (Litviňuková et al. [28]). For mouse, the gastrulation and early-organogenesis atlas was obtained from ArrayExpress (E-MTAB-6967) [30], the Tabula Muris Senis Smart-seq2 dataset from figshare (https://doi.org/10.6084/m9.figshare.8273102; GEO GSE132042) [29], and the SPLiT-seq developing brain and spinal cord dataset from GEO (GSE110823) [31]. Tumor single-cell datasets used for genomic-state inference were obtained from GEO: head and neck squamous cell carcinoma (GSE103322) [37], glioblastoma (GSE131928; Broad Single Cell Portal SCP393) [38] and melanoma (GSE72056) [39]. Spatial-transcriptomics benchmarks used 10x Visium datasets distributed by 10x Genomics. Cross-domain (non-biological) benchmark inputs for astronomy, seismology, remote sensing, climate science, fluid dynamics and molecular dynamics are enumerated with their original sources in Supplementary Data 4. To make the exact file consumed by each benchmark tier reproducible, every benchmark input including derived size tiers, subsamples and preprocessed Scanpy and Seurat checkpoints is mirrored in the project’s Hugging Face dataset repository (https://huggingface.co/datasets/elliotxie/autozyme-datasets). Per-dataset provenance (organism, source and accession, primary publication, license and mirror path) and per-task usage (task × tier × dataset) are provided in Supplementary Data 4 and Supplementary Note 1.

## Code availability

AutoZyme is publicly available at https://github.com/ElliotXie/autozyme under the MIT License and will be archived at Zenodo with a permanent DOI upon publication. The repository contains both the AutoZyme framework, the zyme command-line agent pipeline that developers run to optimize new functions and AutoZyme-Library, the distribution of validated drop-in optimizations (R and Python packages) that end users install to accelerate existing workflows. An interactive web portal for community requests and prioritization is available at https://www.autozyme.com.

## Acknowledgements

We thank the members of the Kendziorski lab for helpful discussions. This work was supported by R35GM162239 (CK PI).

## Author contributions

E.X. conceived the study, designed and implemented the AutoZyme framework, performed the benchmarking and validation, and designed the figures. L.C., Y.C. and J.S. performed testing and debugging and proofread the manuscript. C.K. and E.X. co-wrote the manuscript; C.K. also provided guidance on validation and scope.

